# Whole-catalog reference distributions for reading one person’s polygenic scores, built by an agent-optimized GPU scorer

**DOI:** 10.64898/2026.09.28.755102

**Authors:** Adam Afyouni, Boyuan Yang, Cong Liu

## Abstract

**Objective:** A polygenic score (PGS) is interpretable only as a percentile within a reference distribution, so a person who wants their risk across many diseases/traits needs every candidate model calibrated against a reference distribution. A precalculated frozen reference scoring every genome in a reference panel against every PGS model requires extensive computing resources using current software.

**Materials and Methods:** We reformulated PGS calculation as a sparse matrix product on a GPU and had an autonomous coding agent optimize it under strict numerical validation against pgsc_calc, once on a frozen synthetic training set and once on a 1000 Genomes (1000G) training set, and reran the synthetic training loop three times to assess the variances.

**Results:** On its synthetic training set the scorer sped up 10.6-fold and ran 71-fold faster than pgsc_calc. On 10 genomes and 10 models from the 1000G validation set, the synthetic-trained scorer was 15.3-fold slower than the 1000G-trained version and diverged from pgsc_calc (maximum absolute difference 3.7×10^−1^ vs. 3.1×10^−4^). Across three reruns, agent trajectory variance explained 91 to 96% of wall-clock variance. A memory-bounded rebuild then scored all 3,202 individuals’ genotypes against all 5,385 models in 6.5 h within a 72 GB memory cap.

**Discussion:** Checking an agent’s code only on the data it was tuned on cannot catch overfitting, and reruns of the same prompt differ widely.

**Conclusion:** We release ancestry-stratified reference distributions for the whole PGS Catalog, so a new person’s score can be read without rescoring the reference panel.

## BACKGROUND AND SIGNIFICANCE

A polygenic score (PGS) summarizes an individual’s genome-wide genetic liability for a trait or disease as a weighted sum, s = Σ w_v_·d_v_, where d_v_ is the dosage of the effect allele at variant v and w_v_ is a weight estimated from genome-wide association studies.[1] Once a genome is genotyped, scoring additional diseases costs only computation, and programs such as eMERGE report multiple validated PGS to participants at once (10 scores, planned for 25,000 participants).[2, 3] A raw PGS is unitless, however. It becomes interpretable only as a percentile or ancestry-adjusted z-score within a reference distribution.[2, 4] Reporting many scores therefore requires scoring every reference genome (N) against every model (M), a sparse N × M matrix multiplication. Both dimensions are large and growing. The PGS Catalog held 5,385 harmonized scoring files on 1 September 2026, up from 4,850 in August 2024, ranging from a single variant to 10.3 million variants each.[4, 5] Reference cohorts such as UK Biobank,[6] All of Us,[7] 1000 Genomes,[8] the Million Veteran Program,[9] FinnGen[10] and BioBank Japan[11] range from thousands to hundreds of thousands of individuals of varied ancestry. Even 1000 Genomes (3,202 genomes[8]) yields 17.2 million genotype–model pairs against the full Catalog. Biobank-scale references multiply this by about two orders of magnitude. Frozen, precomputed references avoid rescoring the panel for each patient, but each is valid only for one variant set and one reference population. eMERGE, for example, calibrated its ten scores on 13,475 All of Us participants genotyped and imputed exactly as its own participants were.[2] A service accepting genomes, exomes and multiple arrays must either impute every person or maintain a reference for each platform and population. Each such reference must also be extended whenever the Catalog grows. The central bottleneck is therefore repeatedly recomputing the full N × M matrix. Making that fast would allow calibrated, person-specific, multi-disease reporting in precision medicine practice.

Current tools do not solve this. The community’s best practice is pgsc_calc, the PGS Catalog’s own calculator. It is a Nextflow pipeline that harmonizes scoring files, matches variants against target genotypes and computes scores through PLINK2.[4, 12] In each run the reference panel is scored again: its variants are first intersected with the target genomes, so that target and reference are scored on the same variants.[4, 13] Most of the time goes to matching variants. Researchers estimate that pgsc_calc needs about 135 hours to score one genome against the whole Catalog.[14] Other tools speed up large calculations, including PRSice-2,[15] PLINK2’s own scoring[16] and Spark-based parallelization.[17] Each of them still prepares one target dataset and then scores it against a set of weights.

GPU acceleration is now routine in alignment, variant calling, association testing and population-genetic statistics.[18–23] Polygenic scoring has received comparatively little attention, though some efforts have tried to accelerate it in the Julia programming language.[24] The most direct GPU effort is G-PGS. It scores one patient against the entire Catalog so that a clinician could read that patient’s risk across many diseases in a single report.[14] G-PGS still follows pgsc_calc’s design for the reference. For each patient it scores the reference panel with PLINK2 on the CPU and caches the scores under a hash of the patient’s variant set. Those scores are reused only when a later patient’s variants match.[14] Scoring one patient against the whole Catalog takes 78 minutes. About 70 of those minutes go to the ancestry step, which is dominated by scoring 3,330 reference genomes.[14] In other words, most of the time spent on a new patient goes to scoring other people’s genomes. Precomputing a frozen reference removes this per-patient cost, and once built it can be reused for every patient whose data match it.

## OBJECTIVE

Our goal is to make building a whole-Catalog reference fast enough to support near-real-time personalized genetic risk reports. To achieve this, we use a self-improving agent based on a large language model (LLM) to iteratively develop and optimize the GPU scoring program. We asked two questions. First, whether reformulating polygenic scoring as one sparse product can reduce the computing time for the cohort-by-catalog cross product, correctly enough to yield reference distributions that can be frozen and reused. Second, how much an agent’s optimization target still holds when its output is evaluated on a different distribution from the one it was optimized on, and how much of the outcome depends on the agent’s own run-to-run variation (Figure 1).

**Figure 1.**
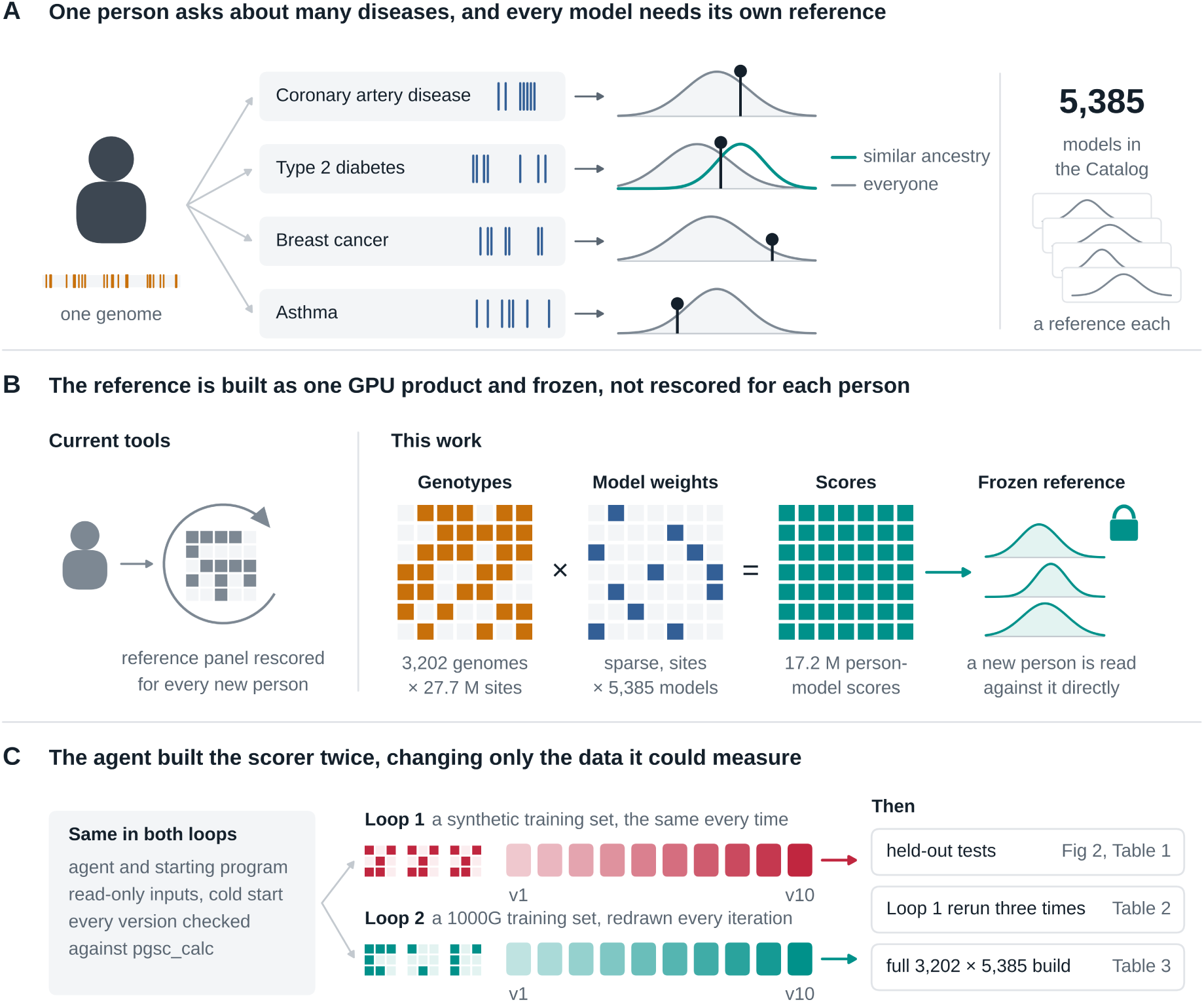
One person, many models, and how the reference behind them is built. (A) A person’s genome can be scored against many published models, and each score is read against that model’s own reference distribution. The curves are illustrations; the second row shows that the same score falls at a different percentile against everyone than against people of similar ancestry. (B) Current tools rescore the reference panel for each new target. Here the panel is scored once as a single sparse matrix product on a GPU, genotypes (3,202 genomes × 27.7 M sites) times sparse model weights (sites × 5,385 models), giving 17.2 million person-model scores whose distributions are frozen so that a new person is read against them directly. (C) The scorer was developed in two ten-version loops by the same agent, from the same starting program and under the same contract, differing only in the data the agent could measure: a frozen synthetic training set (Loop 1, red) or a 1000G training set with genomes and scoring files redrawn at every iteration (Loop 2, teal). The boxes on the right give where each later step is reported.

## MATERIALS AND METHODS

### Data

A synthetic training set of 100 AllofUs-like genotype VCFs, each with 7.65 million variant sites, plus 100 PGS Catalog models randomly drawn, was held read-only throughout the optimization loops and the replicate loops. A synthetic validation set of five shapes (different variant sites and PGS models) was never exposed to any agent loop (Table S1). Real genotypes were drawn from the NYGC 1000 Genomes Project (1000G) high-coverage release,[8] which holds 3,202 individuals and 73,554,796 variant records over chr1 to chr22 and chrX on GRCh38, 1.9 TB in total. PGS models were drawn from the PGS Catalog’s harmonized files, 5,385 models and 3.45 billion rows in total.[4, 5]

### Hardware and baselines

All measurements were single-device, single-host and bare metal, on an NVIDIA GB10 (Grace-Blackwell) GPU with 128 GB of unified memory and a 20-core Arm host CPU. For comparability, every implementation was run with no variant-overlap threshold, no genotype imputation and no ancestry adjustment, except the final reference release described below. Two baselines depend only on existing published software: pgsc_calc v2.3.0 with 20 CPU cores, and a per-individual GPU implementation, a pre-existing hand-written CUDA and CuPy pipeline that converts one genome at a time to a binary store and scores it against all models.[14] No agent read, profiled, edited or optimized the baselines.

### Optimization contract

An autonomous LLM coding agent[25] (Claude Code, Opus-5 model, headless non-interactive mode, with no project-specific configuration and no custom tooling) was given a working cohort-wide GPU scoring script and instructed to make it faster over 10 iterations, measuring and revising the code in a loop like that of self-improving coding agents.[26] It received one architectural hint, to score the whole cohort at once, the domain rules above, and an instruction to keep optimizations general rather than tuned to the specific input’s contents. Each iteration required the agent to (i) profile the previous iteration’s run to identify the bottleneck, (ii) record a written plan, (iii) implement a self-contained script, (iv) execute it from a cold start, and (v) verify the resulting score matrix against a pgsc_calc reference with a hard numerical gate (allclose, atol=1×10^−6^, rtol=1×10^−5^). A speed-up that failed this gate was not considered valid. We did not select which stage to optimize in any iteration, and we did not manually write or debug the GPU kernels the agent produced.

### Two optimization loops and held-out evaluation

The first loop (synthetic training loop) optimized based on the synthetic training set, where pgsc_calc was used to calculate pair-wise scores. The second (1000G training loop) held the contract fixed and varied only the training data: before each iteration, a 1000G training set of 3 to 5 genomes and 3 to 5 PGS Catalog scoring files was drawn randomly, and a fresh pgsc_calc reference was computed for that draw. We call the versions of the two loops synthetic-trained and 1000G-trained. All twenty resulting versions and both baselines were timed on the synthetic validation set and on a 1000G validation set of four shapes, with three repeats each and a 300 s per-run cap on 1000G data. Each 1000G repeat redrew its genomes and models. When a scorer did not finish all three repeats within the cap, its mean is over the repeats that finished, and we report how many finished.

### Replication of the optimization loop

To quantify the stochasticity of the LLM generation process, the synthetic training loop prompt was executed three further times, each as an independent agent process with no shared context and its own workspace. Each replicate’s final version was measured on 1000G validation data at five shapes, with three independent data draws per shape and two executions per draw at the four small shapes, and the 1000G-trained scorer was run on the identical draws as an external reference. Variance was decomposed on log_10_ wall clock with a two-way random-effects model of scorer by data draw, giving components for the agent run, the data draw, their interaction and re-execution.

### Memory-bounded scaling and the final reference build

Memory optimization started from the 1000G-trained final scorer (version 10) and was constrained to memory behavior only. The polygenic score arithmetic and the existing time-reduction strategies were not to change, and the numerical output had to stay identical on any shape both versions could run. Every run was held under a 72 GB host memory cap with swap disabled. The final released reference adds four rules. A locus enters a model’s scoring set S if it matched the model and the panel’s call rate is at least 0.95, with coverage |S|/N reported per model and a deployability flag set below 0.75. Residual missingness inside S is mean-imputed to twice the frozen panel effect-allele frequency. A sample covering less than 0.99 of |S| is excluded from that model’s reference mean and standard deviation. Scores are standardized both pooled and as the standardized residual of a regression on the top ten genotype principal components, with the loadings frozen for projection. Reference statistics use the 2,590 unrelated samples.

### Accuracy reporting

Concordance between each scorer and the corresponding pgsc_calc reference was quantified as the maximum absolute difference across all matrix cells and as Pearson’s r. For the replicates, concordance is reported against the validated 1000G-trained scorer, per model across samples. No formal hypothesis tests were performed.

## RESULTS

### Baselines

pgsc_calc v2.3.0 scored the 100-genotype × 100-model synthetic training set in 1,241 s on 20 CPU cores. Raising the per-process CPU allocation from 2 cores to 20 reduced wall clock only from 1,874 s to 1,241 s, and the dominant stage, the combine step of variant matching, only 1.3-fold (Table S2). The pipeline’s scaling limit lies in matching rather than in scoring. The per-individual GPU implementation took 2,268.5 s on the same set, of which 1,665.0 s was per-sample VCF conversion and 30.0 s the GPU kernel itself (Table S3).

### Optimization on the synthetic training set

Over ten iterations the agent reduced end-to-end wall clock on the synthetic training set from 184.9 s to 17.4 s, a 10.6-fold gain (Figure 2A, Table S4), which is 71-fold faster than the CPU reference and 130-fold faster than the per-individual GPU implementation on the same host. VCF parsing fell from 101.8 s to 13.4 s and model loading plus matching from 76.7 s to 3.4 s, while the sparse product itself fell from 1.9 s to 0.5 s. Accuracy was constant along the trajectory at r = 1.000000 against a non-imputing pgsc_calc reference.

**Figure 2.**
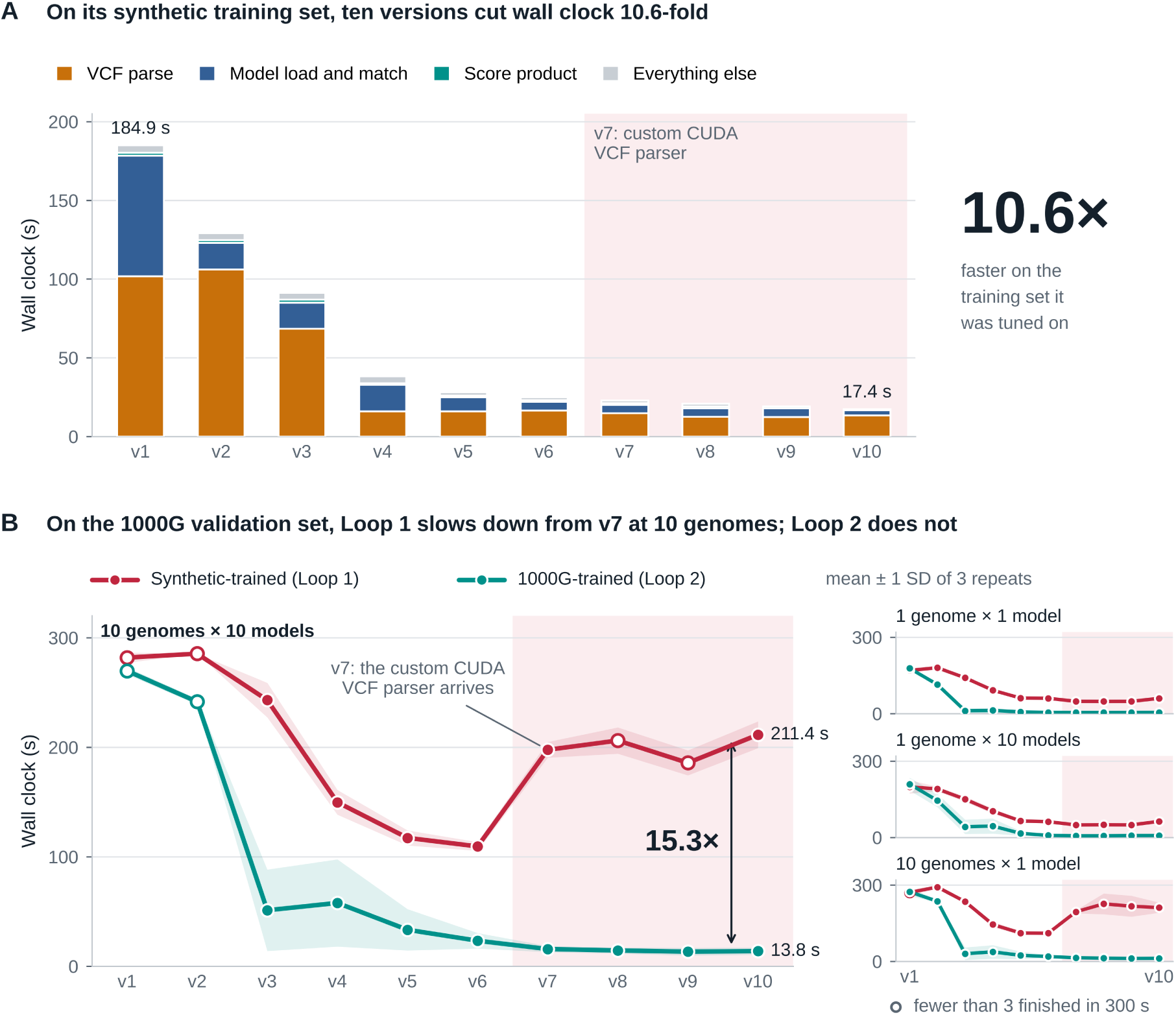
Optimization trajectories of the two loops. (A) Version-by-version wall clock of Loop 1 on its frozen 100 × 100 synthetic training set, split by stage. Everything else is the part of the total not assigned to the three named stages, and each version has one timed run. (B) All ten versions of both loops on the 1000G validation set, at 10 genomes × 10 models (large) and at the three smaller shapes (right). Lines are means of three repeats and bands are ±1 SD; an open marker means fewer than three repeats finished within the 300 s cap, with the mean taken over those that did. The tint marks v7 onward, where Loop 1 introduced a custom CUDA VCF parser; from there its versions get slower at the two 10-genome shapes while its time on the synthetic training set keeps improving. At v10 and 10 × 10 the two loops differ 15.3-fold (211.4 s against 13.8 s).

### Optimization on the resampled 1000G training set

The second loop’s own timings are taken on a redrawn input and are therefore not comparable from iteration to iteration (Table S5), so we assessed it on the four shapes of the 1000G validation set, which the agent was not allowed to see (Figure 2B, Table S6). At 10 genomes × 10 models wall clock fell from 269.7 ± 3.3 s at v1 to 13.8 ± 3.3 s at v10, a 19.5-fold gain, and the gain was 22.8-fold to 36.8-fold at the three other shapes. Every version from v5 onward is faster than every version before v5 at all four shapes, but the last three iterations lie within one standard deviation of each other.

### Held-out validation

All 330 synthetic validation runs completed, and 242 of the 264 runs on the 1000G validation set completed. The failures were pgsc_calc, the per-individual GPU implementation, the earliest agent versions of both families and one 10 × 10 run each of synthetic-trained v8 and v9. On the synthetic validation set the synthetic-trained scorer was fastest, or within noise of fastest, on all five shapes (Table 1). At the largest synthetic shape, 5 genotypes × 100 models, it took 4.06 ± 0.07 s, against 416.4 ± 1.7 s for pgsc_calc and 7.45 ± 0.24 s for the 1000G-trained scorer. On the 1000G validation set the ordering inverted (Table 1, Tables S6 and S7). At 10 genomes × 10 models the four versions fastest on the synthetic training set (v7 to v10), all built around a custom CUDA VCF parser, took 186 to 211 s, which is slower than the mid-trajectory versions v5 and v6 at 109 to 117 s. That parser never learned to discard irrelevant variants, so it pays the full cost of a 73.6-million-variant callset. The 1000G-trained scorer reached 13.8 ± 3.3 s on the same shape, 15.3-fold faster than the synthetic-trained scorer and 21.3-fold faster than pgsc_calc.

**Table 1.**
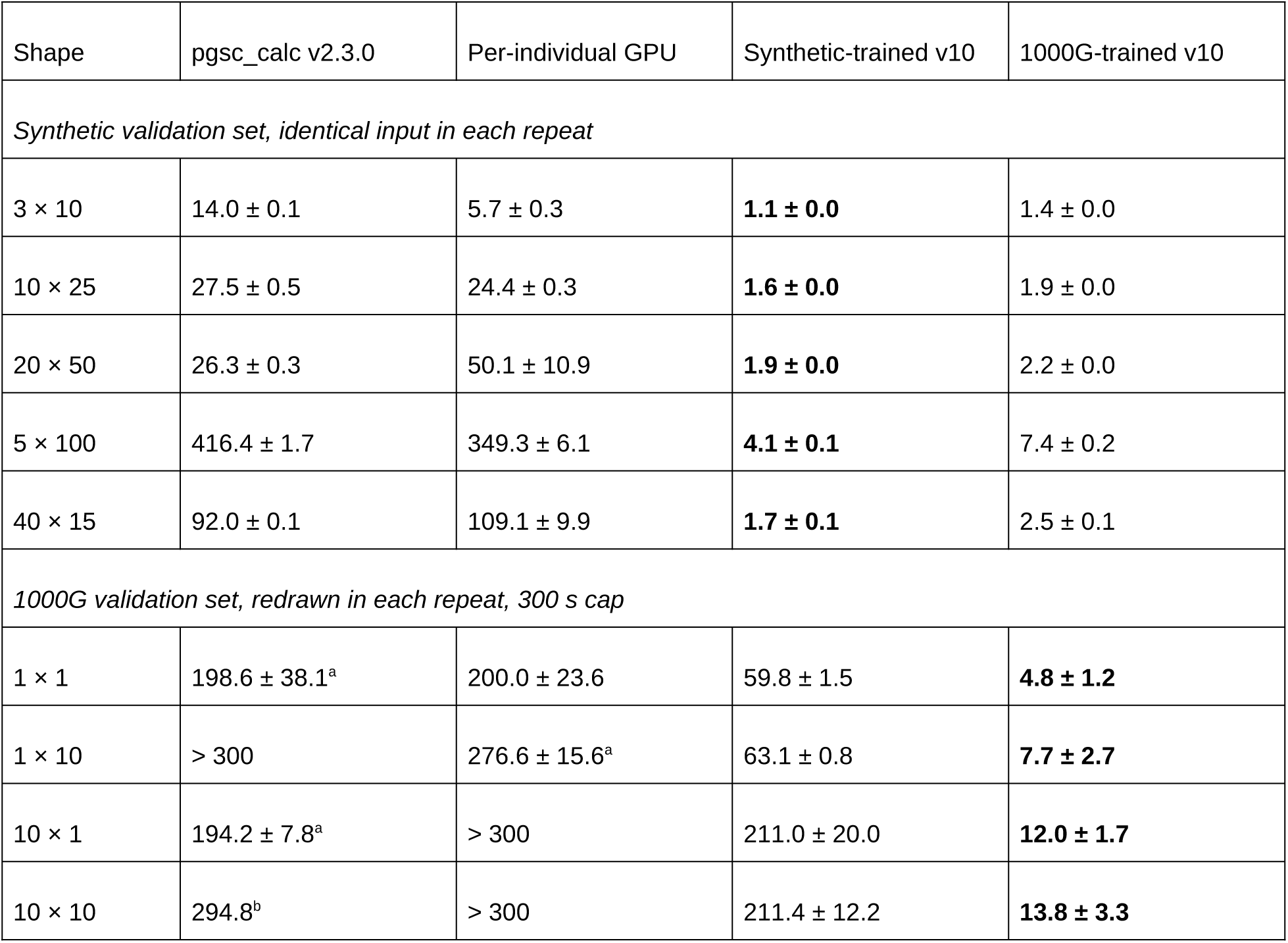
Final scorers on held-out data. Wall clock in seconds, mean ± SD of three repeats, for pgsc_calc v2.3.0, the per-individual GPU implementation and the final version (v10) of each optimization loop. Synthetic shapes repeat the identical input, so their spread is execution jitter alone. 1000G shapes redraw their genomes and models in every repeat and run under a 300 s cap. The fastest scorer in each row is in bold. ^a^Two of three repeats finished within the cap, and the mean is over those two. ^b^One of three finished, so there is no SD. > 300, no repeat finished, so the value is a lower bound and not a measured time. Shapes are genomes × models, and the values are as in Tables S6 and S7.

Concordance with pgsc_calc separated the two families in the same direction. On the synthetic validation set both agreed with pgsc_calc to r = 1.000000 (Table S8). On the 1000G validation set at 10 genomes × 10 models, every synthetic-trained version disagreed by 3.7×10^−1^, while the 1000G-trained family disagreed by 3.1×10^−4^ from v5 onward, three orders of magnitude tighter, a residual dominated by pgsc_calc’s own float32 accumulation (Table S9).

### Run-to-run variance

The coding agent samples its own plan at every iteration, so the artifact it delivers is drawn from the prompt rather than a function of it. The three reruns of the identical prompt produced final programs of comparable size, from 1,025 to 1,118 lines, but of different architectures, with speed-ups on the synthetic training set of 4.2-fold to 17.2-fold (Table 2, Table S10). On 1000G validation data the ranking inverted. The fastest replicate on the synthetic training set, at 7.4 s, was one of two that could not score 100 real genomes at all and failed with a reproducible out-of-memory kill in every repeat. The slowest on the synthetic training set, at 21.8 s, was the only one that completed the target cohort size. At the four small 1000G shapes all three were 2.5-fold to 8.3-fold slower than the 1000G-trained scorer (Tables S11 and S12). A two-way random-effects decomposition of log_10_ wall clock attributes 91.1 to 95.6% of the variance to which agent run produced the scorer, 0.0 to 2.6% to which genomes and models were drawn, and 0.2 to 3.3% to re-executing an identical job (Table S13). The variance is in accuracy as well as wall clock. All three replicates reported a passing correctness check on the synthetic training set, yet on 1000G data one produces scores correlating only 0.834 per model with the validated reference, a typical error of about one standard deviation of that model’s own reference distribution, while the other two stay within about 0.15 standard deviations (Table S14). The mechanisms behind both failures are described in the Supplementary Results.

**Table 2.** Three reruns of the synthetic training loop, timed on 1000G validation data. Replicates A to C are the final programs (v10) of three independent runs of the identical prompt, and the 1000G-trained scorer (v10) is run on the same draws as a reference. Synthetic training is each replicate’s final time on the synthetic training set it optimized (Table S10). The 1000G columns are wall clock in seconds, mean ± SD over three data draws with two executions each, and one execution at 100 × 100 (Table S11). OOM, every run ran out of memory, so no time exists (Table S12). ^a^Five of six runs completed. ^b^Four of six. ^c^Two of three. Median r is the median per-model Pearson correlation with the reference scorer at 10 × 10 (Table S14). In bold, the fastest replicate on the synthetic training set (C) and the one that agrees best with the reference (B).

|  | Synthetic training | 1000G validation data, wall clock (s) |  |  |  |  | Median |
|---|---|---|---|---|---|---|---|
| Scorer | v10 (s) | 1 × 1 | 1 × 10 | 10 × 1 | 10 × 10 | 100 × 100 | r |
| Replicate A | 13.5 | 39.2 ± 0.4 | 41.9 ± 1.2 | 66.4 ± 7.7 | 74.1 ± 7.0<sup>a</sup> | OOM | 0.834 |
| Replicate B | 21.8 | 14.6 ± 1.6 | 16.3 ± 0.5 | 28.0 ± 3.4 | 35.0 ± 6.4 | 160.4 ± 15.5 | **0.999** |
| Replicate C | **7.4** | 35.9 ± 6.0 | 36.0 ± 6.2 | 52.0 ± 1.2<sup>b</sup> | 52.0 ± 1.8<sup>b</sup> | OOM | 0.996 |
| 1000G-trained v10<br>(reference) | n/a | 4.7 ± 0.5 | 5.5 ± 0.6 | 11.2 ± 0.4 | 14.1 ± 2.8 | 137.1 ± 19.8<sup>c</sup> | n/a |

### Memory-bounded scaling to the full cross product

The 1000G-trained scorer still assumes that every structure of the size of the cohort-by-catalog product fits in memory, and it fails at 1,000 genotypes × 1,000 models. Rebuilt inside a fixed memory budget, it completed every run of a shape ladder up to the entire cross product under a 72 GB host memory cap (Table 3, Table S15). The full run scored 3,202 genomes × 5,385 models, 17.2 million genotype-model pairs over 3.45 billion scoring-file rows and 1.9 TB of compressed sequence, in 23,527 s (6.5 h) at a peak of 70.1 GB of memory. The full-scale calculation also moved the bottleneck. Ingestion is 89 to 92% of wall clock at every rung up to 1,000 × 1,000 and the sparse product 2 to 5%, but at full scale ingestion falls to 66% and the product rises to 26% (Table S16).

**Table 3.** Memory-bounded scaling of the released scorer. Every run was held under a 72 GB host-memory cap (Table S15), and the 100 × 100 rung was run three times, with each repeat listed. Ingestion is the two streamed VCF passes, GPU product the cuSPARSE sparse product, and both are given as percentages of wall clock (Table S16). Up to 1,000 × 1,000 ingestion is 89 to 92% of the run and the GPU product 2 to 5%. At the full cross product ingestion falls to 66% and the product rises to 26%.

|  |  |  | Peak host | Share of wall clock (%) |  |  |
|---|---|---|---|---|---|---|
| Genomes × models | Pairs | Wall clock (s) | memory (GB) | Ingestion | GPU product | Other |
| 100 × 100 | 10,000 | 171.5<br>197.1<br>192.8 | 35.3<br>41.0<br>36.8 | 92 | 2 | 6 |
| 500 × 500 | 250,000 | 1,461.4 | 60.1 | 89 | 4 | 7 |
| 1,000 × 1,000 | 1,000,000 | 3,471.1 | 63.6 | 89 | 5 | 6 |
| **3,202 × 5,385** | 17,242,770 | 23,526.6 (6.5 h) | 70.1 | 66 | 26 | 7 |

Building this stage also exposed a defect that every earlier version had missed. cuSPARSE[27] returns an F-contiguous matrix,[28] so the flattening view that the non-additive (dominant and recessive) kernel writes into is a copy rather than a view, and every atomic addition it makes is discarded. Only 4 of the 5,385 catalog models contain a non-additive row, contributing 21 rows, of which 16 matched a target in the full run, 16 out of 3.17 billion matched entries (Table S17).

### Independent validation and the released reference

We recomputed ten randomly selected model-by-genotype combinations from the released cross product with pgsc_calc v2.3.0 under the same configuration. All ten scores agreed, with absolute differences from 0.00000001 to 0.00004619 and relative differences from 1.2e-07 to 4.7e-06, and ten randomly sampled NULL scores produced no score with pgsc_calc either.

The release covers all 3,202 individuals × 5,385 models at 27,683,082 target loci and took 24,052 s (6.7 h) end to end on one GPU and 20 CPU cores (Table S18). Median model coverage is 0.943, and 5,139 of 5,385 models exceed the 0.75 coverage threshold. The remaining 246 models, including the 21 that match no genotypes on the panel, are flagged as not deployable (Table S19). Reference means and standard deviations are computed over the 2,590 unrelated samples, and the 612 trio children are scored but held out. Across the 5,364 models with a defined distribution, the most and the least extreme super-population means are separated by a median of 1.11 pooled standard deviations, by more than one standard deviation for 58.5% of models and by more than two for 12.3%. The top ten genotype principal components explain a median 19.1% of score variance, and more than half of it for 15.7% of models. For a person scored on one of those 15.7% of models, a percentile read against the whole panel mostly reflects their genetic ancestry. Both standardizations are released together with the per-super-population means and the per-model distributions.

## DISCUSSION

We reformulated cohort-scale polygenic scoring as a sparse matrix product on a GPU and delegated its optimization to an off-the-shelf autonomous coding agent under an explicit correctness contract, a setting that KernelBench established for LLM-generated GPU kernels[29] and that related work applies to full research workflows, mathematical discovery and software engineering.[30–32] On its synthetic training set the synthetic-trained scorer was 71-fold faster than the community-standard CPU pipeline, and on the 1000G validation set the 1000G-trained scorer was 21.3-fold faster at 10 genomes × 10 models. GPU scoring is not the bottleneck in small calculations, but at the full 3,202 × 5,385 cross product it became material, at 26% against 66% for VCF ingestion, which locates the scale at which further kernel work would begin to pay.

The central methodological finding is that a correctness gate evaluated only on the distribution used for optimization cannot detect “overfitting” to that distribution. A custom CUDA VCF parser was a clear win on a synthetic training set whose variant sites were generated to overlap the scoring models, and a liability on a real VCF file containing far more sites than any model needs, because nothing in the synthetic training set rewarded discarding irrelevant data. When we held the agent, the underlying model, the contract, the hardware and the starting script fixed, and changed only the data the agent was allowed to train on, the final artifact moved from 211.4 ± 12.2 s to 13.8 ± 3.3 s on the 1000G validation set at 10 genomes × 10 models, and its agreement with the reference implementation improved by three orders of magnitude, because real callsets exercise variant-matching edge cases, such as multi-allelic pooling, sex-chromosome handling and duplicate identifiers, that a synthetic training set generated by the same code path never exercises. An explicit instruction to keep optimizations generalizable did not prevent this, and the overfitting was a correctness problem as well as a speed problem, the correctness problem being the more dangerous of the two. The concern is analogous to overfitting in reinforcement learning[33] or instruction tuning,[34] except that agent self-improvement often lacks a mathematically defined objective. Selecting the shipped version by held-out performance would recover up to about half of the wall-clock loss at negligible additional cost but none of the accuracy loss, and sampling the deployment distribution directly recovers essentially all of both, at the price of a fresh pgsc_calc reference for every 1000G training draw.

A second finding follows from running the same prompt more than once. A single agent run is one sample from a stochastic generation process, and the agent’s run accounted for 91 to 96% of the wall-clock variance across the resulting scorers, which matches recent evidence that run-to-run variance is intrinsic to LLM agents.[35–37] The variance extends to validity. Two of the three replicates could not run the target cohort at all, and one of those two also produced scores correlating only 0.83 per model with a validated reference while its own gate reported success throughout. Optimizing agents are known to satisfy their evaluator without solving the task,[38] kernels that passed KernelBench verification have reported spurious 50- to 120-fold speed-ups by exploiting benchmark loopholes,[39] and flawed task setup or reward design can misestimate agent performance by up to 100%.[40] Variance between agent runs is therefore largely variance in which unrepresentative features each run chose to exploit, so reducing it and removing dependence on a proxy training set are, on this task, the same intervention.

Our release has three components: a lightweight agent optimization kit, which we validated with Claude Code on several hardware configurations including laptops; the final cohort-level scorer; and the reference score distributions, which can be used to compute risk percentiles for new individual scores. One limitation of the reference is that coverage is measured by variant count, so a model can pass the 0.75 threshold while missing some of its highest-weight variants, whose absence affects the score disproportionately.[41] Scoring every model at once is more computation than a single question about a single condition requires, but it is what a growing class of programs does. The eMERGE network returns a genome-informed risk assessment covering eleven chronic diseases to a planned 25,000 participants across ten sites,[3] and for that use the marginal cost of one more condition is one more column rather than one more run.

## CONCLUSION

A frozen, ancestry-stratified reference distribution for each of the 5,364 PGS Catalog models that match the panel lets one person’s scores from many models be read without rescoring the reference panel, and the full cross product ran in 6.5 h on one GPU within a 72 GB memory cap, so it can be repeated for each variant set and reference population. We recommend the released reference for users who want approximate PGS percentiles across many conditions, and the agentic kit or the released scorer for building a custom reference panel where the application needs one.

## ACKNOWLEDGMENTS

We thank Kamalakkannan Ravi, Sanjana Gorlla, and Yundan Zhang for testing the final released agentic optimization kit on their personal laptops. An autonomous coding agent (Claude Code Opus-5) generated the optimized scoring implementations that are the subject of this study, as described in Materials and Methods. AI-assisted technologies were also used during preparation of this manuscript, for language revision and for drafting the figure scripts. The authors reviewed and edited all AI-generated output and took full responsibility for the final content.

## AUTHOR CONTRIBUTIONS

A.A. reproduced the optimization loop on independent hardware, performed the independent validation of the released scores, prepared the figures, and drafted and revised the manuscript; B.Y. wrote the initial agent prompts and markdown files, and ran the repeated analysis; C.L. conceived the study, implemented the scoring pipelines and the release build, ran the optimization loops and replicates, and drafted and revised the manuscript.

## ETHICS

This study used only publicly available, de-identified data from the 1000 Genomes Project and the PGS Catalog. No institutional review board approval or informed consent was required.

## DATA AVAILABILITY

The final scorer, the optimization-loop kit with its prompts, and the audit and validation harness are available at https://github.com/stormliucong/agent_optimized_pgs_calculation. Genotypes are from the NYGC 1000 Genomes Project high-coverage release, and scoring files are from the PGS Catalog,[4, 5] downloaded on 2026-09-01. The reference release itself is distributed as a single bundle with the build parameters, timings and thresholds recorded alongside it. The bundle is deposited at Zenodo (DOI: 10.5281/zenodo.22906325). It contains the frozen scoring manifest, per-locus frozen effect-allele frequencies, PC loadings, panel sample and target tables, and the raw, pooled-standardized, PC-adjusted and per-sample-coverage matrices for all 3,202 individuals × 5,385 models. Note, it should not be applied to variant-only per-sample VCFs without a callable-region definition, because there a locus absent from a sample’s file is ambiguous between hom-ref and uncalled, and the frozen imputation would then be applied to genotypes that were in fact observed.

## SUPPLEMENTARY MATERIAL

Supplementary Methods, Supplementary Results and Tables S1 to S19 are available online.

