## Supplementary Material for "Whole-catalog reference distributions for reading one person’s polygenic scores, built by an agent-optimized GPU scorer"

### Supplementary Methods

#### Datasets

A frozen synthetic training set of 100 AllofUs-like genotype VCFs, each with 7.65 million variant sites, scored against 100 PGS Catalog models, was held read-only throughout optimization loops and throughout the three replicate loops. A synthetic validation set of five shapes (3 to 40 genotypes $\times$ 10 to 100 models, 121 to 366,676 variant sites, and 5.4% to 100% of each shape's model rows carrying a genotype) was never exposed to any agent loop, and its composition is given in Table S1. Real genotypes were drawn from the NYGC 1000 Genomes Project (1000G) high-coverage release,[1] which holds 3,202 individuals, 73,554,796 variant records and 627 to 634 MB compressed per genome, over chr1 to chr22 and chrX on GRCh38, 1.9 TB in total.[1] PGS models were drawn from the PGS Catalog's harmonized files, 5,385 models at time of download, 1 to 10.3 million variants each, median 25,857, 3.45 billion rows and roughly 60 GB in total.[2, 3]

#### Hardware and software

All measurements were single-device, single-host and bare metal, with no cluster, no container and no network filesystem in the I/O path. The host carried an NVIDIA GB10 (Grace-Blackwell) GPU with 128 GB of unified memory shared with the host, a 20-core Arm host CPU (10 performance and 10 efficiency cores), and a 3.6 TB local NVMe SSD holding every input and output. The software stack was CUDA 13.0 with driver 580.95.05, CuPy 14.1.1 with cuSPARSE 12.6.3, ISA-L 2.31.1 for decompression, and Python 3.13, NumPy 2.4, pandas 3.0 and SciPy 1.18 on Ubuntu 24.04. pgsc_calc v2.3.0 was run with the conda profile and 20 CPU cores. For comparability, every implementation was run with no variant-overlap threshold, no genotype imputation and no ancestry adjustment. The one exception is the reference distribution release, where imputation and ancestry adjustment are the product being released and are described separately below.

#### Baseline implementations

Two implementations other than the agent-produced scorers were included as the baselines, which depend only on existing published software, available at `https://github.com/stormliucong/agent_optimized_pgs_calculation`. The first is pgsc_calc v2.3.0 itself, run as described above. On the 100-genotype $\times$ 100-model synthetic training set, it runs the identical pipeline twice with only the per-process CPU allocation changed. The first configuration allocates two CPUs to each heavy Nextflow process, and the second configuration raises that allocation to twenty. Every other pgsc_calc measurement in this study uses the 20-CPU configuration.

The second is the per-individual GPU implementation. It is a pre-existing, hand-written CUDA and CuPy pipeline [4] built on the conventional execution model (`https://github.com/Phenome-Longevity/GPGS`). One individual's VCF is converted to a binary store keyed by a 64-bit variant key made of chromosome, position and 14-bit FNV-1a hashes of REF and ALT, holding float32 dosages. The scoring files are matched against that store by an eight-strategy allele search that excludes strand-ambiguous variants. A per-sample CUDA kernel then produces that individual's scores against all models, using binary search, atomicCAS de-duplication, strand flip, inheritance model, weighted sum and float32 accumulation, with a vectorized rsID lookup recovering rows that the position-based search misses. Scoring N individuals means running the whole pipeline N times, so ingestion is paid once per genome rather than once per cohort. Those design choices, in particular the truncated allele hashes and the float32 accumulation, are why its scores are the least concordant of any implementation reported here. It was wired to each dataset unmodified, with ancestry adjustment disabled so that it computes the same quantity as everything else reported here.

No agent read, profiled, edited or optimized the baseline implementations, and neither is the starting point of either optimization loop.

#### Optimization contract

An autonomous coding agent (Claude Code, Opus-5 model, headless non-interactive mode, no project-specific configuration and no custom tooling beyond its default file, shell and search tools) was given a working cohort-wide GPU scoring script (the starting script had been written by Claude in an interactive browser session from a human's prompt, and its results had not been validated), and instructed to make it faster over ten iterations. The full prompts and the agent's standing instructions are in the agent_optimization_loop folder of the GitHub repository `https://github.com/stormliucong/agent_optimized_pgs_calculation`. It received one architectural hint, which was to score the whole cohort at once, the applicable domain rules of no overlap threshold, no imputation and no ancestry adjustment, and an instruction to keep optimizations general rather than tuned to the specific input's contents. Each iteration required the agent to (i) profile the previous iteration's run to identify the current bottleneck, (ii) record a written plan naming the bottleneck, the evidence for it and the proposed change, (iii) implement a self-contained script for that iteration only, with no code shared across iterations, (iv) execute the script from a cold start, with any temporary caches deleted before the iteration ended and never shared across iterations, and (v) verify the resulting score matrix against a pgsc_calc reference using a hard numerical gate (allclose, atol=$1 \times 10^{-6}$, rtol=$1 \times 10^{-5}$), with the outcome recorded in a run log. A speed-up that failed this gate was not considered valid. We did not select which stage to optimize in any iteration, and we did not manually write or debug the GPU kernels the agent produced.

#### Two optimization loops

The first loop (synthetic training loop) optimized against the frozen synthetic training set described above. The second (1000G training loop) held this contract fixed and varied only the training data. Before each iteration, a 1000G training set of 3 to 5 genomes and 3 to 5 PGS Catalog scoring files was drawn uniformly at random and provided to the agent, and a fresh pgsc_calc reference was computed for that iteration's specific draw, which costs roughly 4 to 6 minutes per reference.

#### Repeated measurement

Every reported wall clock is the mean of three independent repeats, given as mean ± standard deviation. On the fixed synthetic validation shapes all three repeats used the identical input, so the resulting spread reflects execution jitter alone. On the 1000G validation shapes each repeat independently redrew its genomes and models at the stated shape, so the spread reflects both execution jitter and a different dataset instance. Where a scorer did not finish within a 300 s wall-clock cap, the mean is taken over completed repeats only, and the number of completed repeats is reported with it.

#### Replication of the optimization loop

To quantify the stochasticity involved in the LLM generation process, the fixed synthetic training loop prompt was executed three further times, with all three replicates optimized against the frozen synthetic training set. Each replicate ran as an independent agent process with no shared context and no communication with the others, wrote into its own workspace, and was permitted to read only the task description, the starting script, the frozen synthetic training set and the pgsc_calc reference. Each was explicitly instructed to ignore any recalled project context and not to inspect any existing solution. Isolation was by directory and by instruction rather than enforced by the filesystem. Therefore the variance reported is the lower bound in this study. The three ran sequentially and never concurrently, because three jobs sharing one host would corrupt the wall clock each agent optimizes against. Agent optimization loop wall time was 78, 72 and 103 minutes with trajectory details shown in Table S10.

Each replicate's final version (version 10) was then measured on 1000G validation data at five shapes ($1 \times 1$, $1 \times 10$, $10 \times 1$, $10 \times 10$ and $100 \times 100$ genotypes $\times$ models), with three independent data draws per shape and two executions per draw for the four small shapes. The largest shape was run once per draw, because a pass costs about seven minutes and two of four scorers die in it. The previously validated 1000G-trained scorer was run on the identical draws as an external reference. Draws are reproducible and identical across replicates by fixing the random seeds. Replicate order within each shape and draw was rotated so that no replicate sat in the cold-page-cache slot more often than the others. Out-of-memory kills were recorded as results, with the time survived and the kill's origin, either host cgroup or the GPU allocator, rather than retried. This yielded 108 measured runs, 81 from the three replicates and 27 from the reference scorer, of which 96 completed, 12 were killed out of memory and none timed out.

Variance was decomposed on $\log_{10}$ wall clock and the model is a two-way random-effects decomposition of scorer by data draw with two executions per cell, giving components for the agent run, the data draw, their interaction and residual re-execution. Shapes at which a scorer failed with out-of-memory were estimated on the largest complete sub-design. The $100 \times 100$ shape is not estimable, because only one of the three scorers completes it, and it is reported as such.

#### Memory-bounded scaling

Memory optimization started from the 1000G-trained final scorer (v10) and was constrained to memory behaviour only. The polygenic score arithmetic and the existing time-reduction strategies were not to change, and the numerical output had to stay identical on any shape both versions could run. The per-sample accumulators that retained every sample's keys, dosages and ordinals until the last VCF was parsed, about 470 GB at $1{,}000 \times 1{,}000$, were replaced by two streamed VCF passes. The first pass builds the target union and the per-target ordinal incrementally, and the second folds each sample directly into its column of a memory-mapped dosage matrix. Scoring files, 3.45 billion rows and roughly 350 GB of decompressed text in full, are read in budget-sized blocks after a locus-only first pass that decodes contig/chromosome and position. The per-sample member scan walks a 32 MB window through an mmap instead of loading a 628 MB VCF per worker. The scoring tile width is derived from the budget (as an input argument) rather than fixed. The dosage matrix itself, roughly 190 GB for the whole cross product, is memory-mapped so that reclaimable page cache absorbs what RAM cannot. Every run was placed with a 72 GB host memory cap with swap disabled, and an external watchdog that killed the job if machine-wide available memory stayed below 12 GB. Peak memory is reported as total memory minus the minimum machine-wide available memory, sampled twice a second across the whole process tree.

##### Final reference build

In addition, we included four calculation rules for the final release of full cross-product reference. (1) A locus enters a model's scoring set S if it matched the model and the panel's call rate over the reference samples is at least 0.95, with coverage $|S|/N$ reported per model and a deployability flag set below 0.75. (2) Residual missingness inside S is mean-imputed to twice the frozen panel effect-allele frequency, implemented as one additional sparse product on the same sparsity pattern, and non-additive loci are imputed with the expectation of the corresponding transform under Hardy-Weinberg. That frequency is stored per locus in effect-allele orientation, together with the weight, the flip flag and the inheritance model, so that a future individual is imputed with the panel's number; (3) Per-sample coverage is computed as a second product against the binary pattern of the weight matrix, and a sample below 0.99 of $|S|$ is excluded from that model's reference mean and standard deviation. (4) Standardization is applied last, both pooled and as the standardized residual of a regression on the top ten genotype principal components, computed here from common, well-called, physically thinned variants (72,437 variants at 20,000 bp spacing, minor allele frequency at least 0.05, adequate for global-ancestry axes and nothing else) with the loadings frozen for projection. The final reference scores use the 2,590 unrelated samples (612 trio children are scored but excluded from the reference set). A subset of 20 final release scores was independently checked using pgsc_calc under the same configuration rules as described above.

##### Statistical and accuracy reporting

Concordance between each scorer and the corresponding pgsc_calc reference was quantified as the maximum absolute difference across all matrix cells and as Pearson's r. For the replicate comparison, where no pgsc_calc reference was affordable at the shapes measured, concordance is reported against the validated 1000G-trained scorer and computed per model column across samples, because models differ in scale by orders of magnitude and a correlation pooled over the whole matrix would mostly measure scale agreement. No formal hypothesis tests were performed.

#### Supplementary Results

##### How the replicates failed on 1000G data

One replicate builds its cohort variant axis from the VCF union of 73,554,796 sites rather than from the model loci, which is free on a synthetic training set whose VCFs contain only variants the models mention and is about 7.4 GB of int8 at 100 real genomes. The other sizes its axis correctly but admits later VCFs to a fast path only through an exact byte-diff against the first file, and real per-individual 1000 Genomes VCFs are not byte-identical. Its own log reports reusing the first file's site table for six of the ten per-sample files at the ten-genome shape, and the same fallback rate at 100 genotypes puts 20 concurrent workers each doing a full parse of a 628 MB VCF, which is what the kernel kills at a reproducible 62 to 82 s into every attempt. The same two mechanisms recur at smaller shapes as data-driven and execution-driven variance, with one replicate surviving two data draws at $10 \times 10$ and dying on the third in both executions, and another dying on one execution of a draw and completing the identical input on the next. The variance is in accuracy as well as wall clock. All three replicates reported a passing correctness check on the synthetic training set, yet on 1000G data one produces scores correlating only 0.834 per model with the validated reference, with a worst model r of 0.251 and a typical error of about one standard deviation of that model's own reference distribution, while the other two stay within about 0.15 standard deviations (Table S14).

#### Supplementary Tables

**Table S1.** Composition of the five synthetic validation shapes. None was exposed to any agent optimization loop. The shapes vary along four axes independently, namely number of genotypes, number of models, model size, and how much of each model the genotypes actually cover. Shape val 5 additionally contains about 270,000 variants that no model mentions, so its target callset is much wider than its models, which is the axis on which the synthetic-trained scorers later failed on 1000G data.

| Shape | Genotypes | Models | Variant sites | Model rows | Model rows genotyped | Design |
|---|---|---|---|---|---|---|
| val 1 | 3 | 10 | 121 | 139 | 100.0% | tiny, 10 smallest models, every model variant genotyped |
| val 2 | 10 | 25 | 221,120 | 372,046 | 100.0% | small/mid models (<= 50 k variants), full overlap |
| val 3 | 20 | 50 | 183,272 | 375,436 | 89.2% | 50 mid-sized models, 20 k variants sampled per model |
| val 4 | 5 | 100 | 325,000 | 34,800,352 | 5.4% | few samples x 100 models incl. the 6.9 M-variant genome-wide ones, union subsampled so most model rows have no genotype |
| val 5 | 40 | 15 | 366,676 | 2,371,201 | 6.0% | many samples x 15 models, half of each model dropped and ∼270 k model-free variants added, so the VCF is much wider than the models |

**Table S2.** pgsc_calc v2.3.0 core-count scaling on the synthetic training set (100 genotypes × 100 models, 7,653,879 variant sites). The identical pipeline was run twice, changing only the per-process CPU allocation. The stock configuration pins the heavy Nextflow processes to 2 CPUs each, and the second raises that to 20. --max_cpus alone has no effect, being only a ceiling. One run per configuration, and the two produced identical score matrices (maximum absolute difference 0.0). Stage times come from Nextflow's own task trace.

| Stage | Nextflow process | 2 CPUs/process | 20 CPUs/process | Speed-up |
|---|---|---|---|---|
| Input reconciliation (VCF merge) | n/a | 27.0 s | 27.2 s | 0.99× |
| VCF parsing | PLINK2_VCF | 16.0 s | 15.4 s | 1.04× |
| Model loading | FORMAT_SCOREFILES | 570.0 s | 259.0 s | 2.20× |
| Variant matching | MATCH_VARIANTS | 81.0 s | 26.8 s | 3.02× |
| Variant matching (combine) | MATCH_COMBINE | 1,160.0 s | 879.0 s | 1.32× |
| Score calculation | PLINK2_SCORE ×2 | 48.9 s | 44.4 s | 1.10× |
| Score aggregation | SCORE_AGGREGATE | 27.0 s | 26.7 s | 1.01× |
| Pipeline wall clock | | 1,874 s | 1,241 s | 1.51× |

**Table S3.** Per-individual GPU implementation, stage breakdown on the synthetic training set (100 genotypes $\times$ 100 models). Model loading is paid once for the whole run. Every other stage is paid once per genome, and the totals below are summed over the 100 genomes. rsID fallback and the two GPU rows in italics are nested inside GPU scoring and are not added again.

| **Stage** | **Total (s)** | **Per genome (s)** | **% of run** |
|---|---:|---:|---:|
| Model loading / PGS catalog sync (once) | 320.4 | 3.20 | 14.1% |
| VCF parsing $\rightarrow$ per-sample binary store | 1,665.0 | 16.65 | 73.4% |
| Binary store load | 36.7 | 0.37 | 1.6% |
| Weight-file I/O | 71.1 | 0.71 | 3.1% |
| GPU scoring | 175.2 | 1.75 | 7.7% |
| of which, host$\rightarrow$device transfer | 22.7 | 0.23 | 1.0% |
| of which, CUDA reduction kernel | 30.0 | 0.30 | 1.3% |
| of which, vectorized rsID fallback | 122.5 | 1.23 | 5.4% |
| Total | 2,268.5 | 22.68 | 100.0% |

**Table S4.** Synthetic training loop, per-stage wall clock for all ten iterations (100 genotypes $\times$ 100 models). Each column is that iteration's single cold run, which is the measurement the agent optimized against. Stage names change along the trajectory because the agent restructured the pipeline. Version v9 fuses weight-matrix assembly and the sparse product into one double-precision kernel, and v10 splits target-table publication out of model matching so that model loading can overlap parsing. All values are in seconds.

| **Stage** | **v1** | **v2** | **v3** | **v4** | **v5** | **v6** | **v7** | **v8** | **v9** | **v10** |
|---|---|---|---|---|---|---|---|---|---|---|
| VCF parse + dosage matrix | 101.84 | 106.16 | 68.51 | 16.12 | 16.04 | 16.46 | 14.75 | 12.56 | 12.50 | 13.39 |
| Target index build | 2.35 | 2.58 | 2.35 | 2.34 | 0.01 | 0.01 | 0.01 | 0.01 | 0.01 | n/a |
| Publish target table | n/a | n/a | n/a | n/a | n/a | n/a | n/a | n/a | n/a | 0.04 |
| Model loading + matching | 76.69 | 16.75 | 16.56 | 16.75 | 9.12 | 5.58 | 5.36 | 5.38 | 5.53 | n/a |
| Model matching remainder | n/a | n/a | n/a | n/a | n/a | n/a | n/a | n/a | n/a | 3.36 |
| Weight matrix build | 2.10 | 2.08 | 1.98 | 2.02 | 2.02 | 2.02 | 1.75 | 1.86 | n/a | n/a |
| Sparse product (cuSPARSE) | 1.90 | 1.78 | 1.77 | 1.04 | 1.05 | 1.05 | 1.07 | 1.06 | n/a | n/a |
| Fused float64 scoring kernel | n/a | n/a | n/a | n/a | n/a | n/a | n/a | n/a | 1.18 | 0.51 |
| Write output | 0.00 | 0.00 | 0.00 | 0.00 | 0.00 | 0.00 | 0.00 | 0.00 | 0.00 | 0.00 |
| Total | 184.89 | 129.35 | 91.17 | 38.28 | 28.24 | 25.12 | 22.94 | 20.87 | 19.22 | 17.41 |

**Table S5.** 1000G training loop, every iteration on its own draw and every version that was run on the common draw. These are the loop's own training measurements. The main text does not quote them and reports this loop only on the 1000G validation set, in Fig 2. The input is redrawn before each iteration, so the own-draw column is not comparable across rows and is given only to show what each iteration measured. The common draw is iteration 5's draw (5 genomes $\times$ 5 models, 11.66 million model rows), the largest drawn in the loop. Versions v1 to v4 predate it and were never run on it. Gate is the hard numerical check against that iteration's own freshly computed pgsc_calc reference (allclose, atol=$1 \times 10^{-6}$, rtol=$1 \times 10^{-5}$). On the common draw every version listed produced identical target counts, identical matched counts per model and identical scores, so the differences there are scheduling only.

| Version | Own draw, total | Common draw, VCF parse | Common draw, matching | Common draw, total | Gate | Dominant change |
|---|---:|---:|---:|---|---|---|
| v1 | 215.58 s | n/a | n/a | not run | PASSED | Baseline port onto real 1000 Genomes data |
| v2 | 162.87 s | n/a | n/a | not run | PASSED | One parallel task per gzip member |
| v3 | 10.77 s | n/a | n/a | not run | PASSED | Model-driven position prefilter (the loop's largest win) |
| v4 | 19.40 s | n/a | n/a | not run | FAILED (matching rule) | CPU-vectorized VCF parse (negative result, gate failed) |
| v5 | 33.61 s | 22.80 s | 2.27 s | 33.61 s | PASSED | Merged candidate pool fixed, model side off the interpreter |
| v6 | 8.54 s | 13.18 s | 1.46 s | 18.35 s | PASSED | Set algebra on the GPU, dosage matrix stays resident |
| v7 | 8.88 s | 7.15 s | 1.48 s | 12.39 s | PASSED | VCF records parsed by two fused CUDA kernels |
| v8 | 7.43 s | 6.50 s | 1.51 s | 11.63 s | PASSED | One sweep of the VCF text, remaining work overlapped |
| v9 | 6.72 s | 6.00 s | 1.10 s | 10.62 s | PASSED | Decompression moved into worker processes, device pre-pass |
| v10 | 7.26 s | 5.97 s | 1.11 s | 10.55 s | PASSED | Device pipeline batched across gzip members |

**Table S6.** 1000G validation set wall clock (seconds, mean ± sd over 3 repeats), under a 300 s per-run cap. Each repeat independently redraws its genomes and models at the stated shape, so the spread reflects both execution jitter and a different problem instance. Runs killed at the cap are counted rather than dropped, and where all three repeats were killed the entry is the cap. In total 242 of 264 runs completed.

| **Scorer** | **1 × 1** | **1 × 10** | **10 × 1** | **10 × 10** |
|---|---|---|---|---|
| pgsc_calc v2.3.0 (CPU, 20 cores) | 198.6 ± 38.1 [1 capped] | > 300 s (3/3) | 194.2 ± 7.8 [1 capped] | 294.8 [2 capped] |
| Per-individual GPU implementation | 200.0 ± 23.6 | 276.6 ± 15.6 [1 capped] | > 300 s (3/3) | > 300 s (3/3) |
| Synthetic-trained v1 | 169.4 ± 4.8 | 198.8 ± 26.3 | 270.4 ± 10.4 [1 capped] | 281.8 ± 4.7 [1 capped] |
| Synthetic-trained v2 | 180.1 ± 5.6 | 190.5 ± 9.3 | 291.7 ± 7.8 | 285.4 [2 capped] |
| Synthetic-trained v3 | 140.5 ± 7.1 | 150.3 ± 10.8 | 234.5 ± 7.1 | 243.0 ± 15.8 |
| Synthetic-trained v4 | 91.6 ± 5.8 | 103.4 ± 10.9 | 144.6 ± 10.7 | 149.7 ± 11.3 |
| Synthetic-trained v5 | 60.8 ± 3.0 | 64.9 ± 4.8 | 111.3 ± 7.7 | 117.2 ± 6.9 |
| Synthetic-trained v6 | 60.0 ± 3.0 | 61.9 ± 2.3 | 110.6 ± 5.2 | 109.4 ± 4.0 |
| Synthetic-trained v7 | 48.0 ± 2.6 | 49.2 ± 1.8 | 194.6 ± 5.9 | 197.6 ± 7.2 |
| Synthetic-trained v8 | 47.6 ± 2.8 | 49.7 ± 1.9 | 225.8 ± 40.8 | 206.1 ± 12.0 [1 capped] |
| Synthetic-trained v9 | 47.9 ± 2.6 | 49.3 ± 1.3 | 216.6 ± 41.1 | 185.7 ± 11.4 [1 capped] |
| Synthetic-trained v10 | 59.8 ± 1.5 | 63.1 ± 0.8 | 211.0 ± 20.0 | 211.4 ± 12.2 |
| 1000G-trained v1 | 177.7 ± 5.3 | 209.4 ± 25.2 | 273.0 ± 15.2 | 269.7 ± 3.3 [1 capped] |
| 1000G-trained v2 | 114.0 ± 7.4 | 145.1 ± 26.1 | 236.0 ± 6.3 | 241.5 ± 2.7 [1 capped] |
| 1000G-trained v3 | 10.9 ± 9.6 | 42.0 ± 28.2 | 30.1 ± 24.7 | 51.1 ± 37.2 |
| 1000G-trained v4 | 12.3 ± 10.8 | 44.8 ± 29.6 | 37.3 ± 26.6 | 57.8 ± 39.8 |
| 1000G-trained v5 | 6.6 ± 4.3 | 16.1 ± 9.2 | 23.6 ± 14.9 | 33.3 ± 18.9 |
| 1000G-trained v6 | 4.7 ± 1.6 | 8.2 ± 3.1 | 19.1 ± 5.9 | 23.3 ± 7.1 |
| 1000G-trained v7 | 4.4 ± 1.2 | 6.6 ± 2.1 | 14.0 ± 1.9 | 15.7 ± 2.8 |
| 1000G-trained v8 | 4.2 ± 1.1 | 6.6 ± 2.5 | 12.5 ± 1.8 | 14.4 ± 2.8 |
| 1000G-trained v9 | 4.5 ± 1.2 | 7.5 ± 2.6 | 11.5 ± 2.2 | 13.3 ± 3.5 |
| 1000G-trained v10 | 4.8 ± 1.2 | 7.7 ± 2.7 | 12.0 ± 1.7 | 13.8 ± 3.3 |

**Table S7.** Synthetic validation wall clock (seconds, mean ± sd over 3 repeats on the identical input). All 330 of 330 runs completed and no run hit the 900 s cap. The spread here is execution jitter only, because the input does not change between repeats. Shapes are genotypes × models.

| **Scorer** | **val 1 (3×10)** | **val 2 (10×25)** | **val 3 (20×50)** | **val 4 (5×100)** | **val 5 (40×15)** |
|---|---|---|---|---|---|
| pgsc_calc v2.3.0 (CPU, 20 cores) | 14.0 ± 0.1 | 27.5 ± 0.5 | 26.3 ± 0.3 | 416.4 ± 1.7 | 92.0 ± 0.1 |
| Per-individual GPU implementation | 5.7 ± 0.3 | 24.4 ± 0.3 | 50.1 ± 10.9 | 349.3 ± 6.1 | 109.1 ± 9.9 |
| Synthetic-trained v1 | 0.9 ± 0.0 | 2.2 ± 0.0 | 2.2 ± 0.0 | 72.8 ± 3.1 | 7.9 ± 0.4 |
| Synthetic-trained v2 | 0.9 ± 0.0 | 1.6 ± 0.0 | 1.6 ± 0.0 | 16.9 ± 2.3 | 7.5 ± 0.3 |
| Synthetic-trained v3 | 1.0 ± 0.0 | 1.6 ± 0.0 | 1.6 ± 0.0 | 15.7 ± 0.2 | 6.9 ± 0.1 |
| Synthetic-trained v4 | 1.2 ± 0.0 | 1.6 ± 0.0 | 1.6 ± 0.0 | 15.4 ± 0.3 | 6.0 ± 0.3 |
| Synthetic-trained v5 | 1.2 ± 0.0 | 1.5 ± 0.0 | 1.5 ± 0.0 | 8.7 ± 0.1 | 3.5 ± 0.1 |
| Synthetic-trained v6 | 1.2 ± 0.0 | 1.5 ± 0.0 | 1.6 ± 0.0 | 4.8 ± 0.0 | 2.3 ± 0.0 |
| Synthetic-trained v7 | 1.2 ± 0.0 | 1.7 ± 0.0 | 1.9 ± 0.0 | 4.8 ± 0.1 | 2.4 ± 0.0 |
| Synthetic-trained v8 | 1.2 ± 0.0 | 1.8 ± 0.0 | 2.0 ± 0.0 | 4.9 ± 0.1 | 2.6 ± 0.1 |
| Synthetic-trained v9 | 1.1 ± 0.0 | 1.7 ± 0.0 | 1.9 ± 0.0 | 4.6 ± 0.0 | 2.5 ± 0.0 |
| Synthetic-trained v10 | 1.1 ± 0.0 | 1.6 ± 0.0 | 1.9 ± 0.0 | 4.1 ± 0.1 | 1.7 ± 0.1 |
| 1000G-trained v1 | 0.9 ± 0.0 | 2.3 ± 0.1 | 2.3 ± 0.0 | 77.5 ± 0.4 | 7.9 ± 0.4 |
| 1000G-trained v2 | 0.9 ± 0.0 | 2.4 ± 0.0 | 2.4 ± 0.1 | 77.9 ± 0.3 | 7.9 ± 0.0 |
| 1000G-trained v3 | 0.9 ± 0.0 | 2.2 ± 0.1 | 2.1 ± 0.1 | 67.1 ± 0.4 | 8.2 ± 0.1 |
| 1000G-trained v4 | 0.9 ± 0.0 | 2.3 ± 0.1 | 2.2 ± 0.0 | 67.8 ± 3.6 | 8.9 ± 0.1 |
| 1000G-trained v5 | 1.1 ± 0.0 | 2.0 ± 0.1 | 2.1 ± 0.0 | 14.3 ± 0.4 | 4.4 ± 0.1 |
| 1000G-trained v6 | 1.3 ± 0.0 | 2.1 ± 0.0 | 2.2 ± 0.0 | 8.7 ± 0.2 | 3.3 ± 0.0 |
| 1000G-trained v7 | 1.2 ± 0.0 | 1.7 ± 0.0 | 1.8 ± 0.0 | 6.9 ± 0.2 | 2.2 ± 0.1 |
| 1000G-trained v8 | 1.2 ± 0.0 | 1.7 ± 0.0 | 1.8 ± 0.0 | 7.0 ± 0.4 | 2.3 ± 0.1 |
| 1000G-trained v9 | 1.3 ± 0.0 | 2.4 ± 0.9 | 2.1 ± 0.0 | 7.4 ± 0.2 | 2.4 ± 0.0 |
| 1000G-trained v10 | 1.4 ± 0.0 | 1.9 ± 0.0 | 2.2 ± 0.0 | 7.4 ± 0.2 | 2.5 ± 0.1 |

**Table S8.** Synthetic validation concordance with pgsc_calc, worst-case absolute difference over all matrix cells and all three repeats, with the minimum Pearson r in parentheses. Each run is compared against the pgsc_calc matrix of the same shape and repeat. pgsc_calc is its own reference and is shown as 0 for completeness. The per-individual GPU implementation is two to five orders of magnitude looser than the agent versions, while both agent families agree with pgsc_calc at r = 1.000000 on synthetic data, which is the point made in the main text.

| Scorer | val 1 | val 2 | val 3 | val 4 | val 5 |
|---|---|---|---|---|---|
| pgsc_calc v2.3.0 (CPU, 20 cores) | 0 (1.000000) | 0 (1.000000) | 0 (1.000000) | 0 (1.000000) | 0 (1.000000) |
| Per-individual GPU implementation | 2.00e-01 (0.999745) | 7.59e+00 (0.999998) | 5.25e+00 (0.999983) | 3.61e+00 (0.999992) | 5.08e+00 (0.999985) |
| Synthetic-trained v1 | 4.00e-06 (1.000000) | 2.76e-02 (1.000000) | 1.01e-02 (1.000000) | 1.20e-02 (1.000000) | 2.23e-03 (1.000000) |
| Synthetic-trained v2 | 4.00e-06 (1.000000) | 2.76e-02 (1.000000) | 1.01e-02 (1.000000) | 1.20e-02 (1.000000) | 2.23e-03 (1.000000) |
| Synthetic-trained v3 | 4.00e-06 (1.000000) | 2.76e-02 (1.000000) | 1.01e-02 (1.000000) | 1.20e-02 (1.000000) | 2.23e-03 (1.000000) |
| Synthetic-trained v4 | 4.00e-06 (1.000000) | 2.76e-02 (1.000000) | 1.01e-02 (1.000000) | 1.20e-02 (1.000000) | 2.23e-03 (1.000000) |
| Synthetic-trained v5 | 4.00e-06 (1.000000) | 2.76e-02 (1.000000) | 1.01e-02 (1.000000) | 1.20e-02 (1.000000) | 2.23e-03 (1.000000) |
| Synthetic-trained v6 | 4.00e-06 (1.000000) | 2.76e-02 (1.000000) | 1.01e-02 (1.000000) | 1.20e-02 (1.000000) | 2.23e-03 (1.000000) |
| Synthetic-trained v7 | 4.00e-06 (1.000000) | 2.76e-02 (1.000000) | 1.01e-02 (1.000000) | 1.20e-02 (1.000000) | 2.23e-03 (1.000000) |
| Synthetic-trained v8 | 4.00e-06 (1.000000) | 2.76e-02 (1.000000) | 1.01e-02 (1.000000) | 1.20e-02 (1.000000) | 2.23e-03 (1.000000) |
| Synthetic-trained v9 | 4.00e-06 (1.000000) | 2.76e-02 (1.000000) | 1.01e-02 (1.000000) | 1.20e-02 (1.000000) | 2.23e-03 (1.000000) |
| Synthetic-trained v10 | 4.00e-06 (1.000000) | 2.76e-02 (1.000000) | 1.01e-02 (1.000000) | 1.20e-02 (1.000000) | 2.23e-03 (1.000000) |
| 1000G-trained v1 | 4.00e-06 (1.000000) | 5.00e-04 (1.000000) | 5.00e-04 (1.000000) | 5.00e-04 (1.000000) | 5.00e-04 (1.000000) |
| 1000G-trained v2 | 4.00e-06 (1.000000) | 5.00e-04 (1.000000) | 5.00e-04 (1.000000) | 5.00e-04 (1.000000) | 5.00e-04 (1.000000) |
| 1000G-trained v3 | 4.00e-06 (1.000000) | 5.00e-04 (1.000000) | 5.00e-04 (1.000000) | 5.00e-04 (1.000000) | 5.00e-04 (1.000000) |
| 1000G-trained v4 | 4.00e-06 (1.000000) | 5.00e-04 (1.000000) | 5.00e-04 (1.000000) | 5.00e-04 (1.000000) | 5.00e-04 (1.000000) |
| 1000G-trained v5 | 4.00e-06 (1.000000) | 5.00e-04 (1.000000) | 5.00e-04 (1.000000) | 5.00e-04 (1.000000) | 5.00e-04 (1.000000) |
| 1000G-trained v6 | 4.00e-06 (1.000000) | 5.00e-04 (1.000000) | 5.00e-04 (1.000000) | 5.00e-04 (1.000000) | 5.00e-04 (1.000000) |
| 1000G-trained v7 | 4.00e-06 (1.000000) | 5.00e-04 (1.000000) | 5.00e-04 (1.000000) | 5.00e-04 (1.000000) | 5.00e-04 (1.000000) |
| 1000G-trained v8 | 4.00e-06 (1.000000) | 5.00e-04 (1.000000) | 5.00e-04 (1.000000) | 5.00e-04 (1.000000) | 5.00e-04 (1.000000) |
| 1000G-trained v9 | 4.00e-06 (1.000000) | 5.00e-04 (1.000000) | 5.00e-04 (1.000000) | 5.00e-04 (1.000000) | 5.00e-04 (1.000000) |
| 1000G-trained v10 | 4.00e-06 (1.000000) | 5.00e-04 (1.000000) | 5.00e-04 (1.000000) | 5.00e-04 (1.000000) | 5.00e-04 (1.000000) |

**Table S9.** 1000G validation set concordance with pgsc_calc, worst-case absolute difference over all matrix cells and all repeats in which both the scorer and pgsc_calc completed, with the minimum Pearson r in parentheses. Blank where pgsc_calc or the scorer was capped in every repeat at that shape. At 10 × 10 the synthetic-trained family sits at $3.7 \times 10^{-1}$ and the 1000G-trained family at $3.1 \times 10^{-4}$ from v5 onward, which is the three-orders-of-magnitude separation reported in the main text. The 1000G-trained family's residual disagreement is dominated by pgsc_calc's own float32 accumulation.

| **Scorer** | **1 × 1** | **10 × 1** | **10 × 10** |
|---|---|---|---|
| pgsc_calc v2.3.0 (CPU, 20 cores) | 0 (n/a) | 0 (1.000000) | 0 (1.000000) |
| Per-individual GPU implementation | 1.10e-07 (n/a) | capped in all repeats | capped in all repeats |
| Synthetic-trained v1 | 4.00e-07 (n/a) | 2.00e-06 (1.000000) | 3.68e-01 (0.999999) |
| Synthetic-trained v2 | 4.00e-07 (n/a) | 5.00e-06 (1.000000) | 3.68e-01 (0.999999) |
| Synthetic-trained v3 | 4.00e-07 (n/a) | 3.00e-06 (1.000000) | 3.68e-01 (0.999999) |
| Synthetic-trained v4 | 4.00e-07 (n/a) | 6.00e-06 (1.000000) | 3.68e-01 (0.999999) |
| Synthetic-trained v5 | 4.00e-07 (n/a) | 5.00e-06 (1.000000) | 3.68e-01 (0.999999) |
| Synthetic-trained v6 | 4.00e-07 (n/a) | 1.00e-05 (1.000000) | 3.68e-01 (0.999999) |
| Synthetic-trained v7 | 4.00e-07 (n/a) | 3.00e-06 (1.000000) | 3.68e-01 (0.999999) |
| Synthetic-trained v8 | 4.00e-07 (n/a) | 3.00e-06 (1.000000) | 3.68e-01 (0.999999) |
| Synthetic-trained v9 | 4.00e-07 (n/a) | 5.00e-07 (1.000000) | 3.68e-01 (0.999999) |
| Synthetic-trained v10 | 4.00e-07 (n/a) | 5.00e-07 (1.000000) | 3.68e-01 (0.999999) |
| 1000G-trained v1 | 4.00e-07 (n/a) | 5.00e-07 (1.000000) | 3.28e-03 (1.000000) |
| 1000G-trained v2 | 4.00e-07 (n/a) | 5.00e-07 (1.000000) | 3.28e-03 (1.000000) |
| 1000G-trained v3 | 4.00e-07 (n/a) | 5.00e-07 (1.000000) | 3.28e-03 (1.000000) |
| 1000G-trained v4 | 4.00e-07 (n/a) | 5.00e-07 (1.000000) | 1.34e-03 (1.000000) |
| 1000G-trained v5 | 4.00e-07 (n/a) | 5.00e-07 (1.000000) | 3.07e-04 (1.000000) |
| 1000G-trained v6 | 4.00e-07 (n/a) | 5.00e-07 (1.000000) | 3.07e-04 (1.000000) |
| 1000G-trained v7 | 4.00e-07 (n/a) | 5.00e-07 (1.000000) | 3.07e-04 (1.000000) |
| 1000G-trained v8 | 4.00e-07 (n/a) | 5.00e-07 (1.000000) | 3.07e-04 (1.000000) |
| 1000G-trained v9 | 4.00e-07 (n/a) | 5.00e-07 (1.000000) | 3.07e-04 (1.000000) |
| 1000G-trained v10 | 4.00e-07 (n/a) | 5.00e-07 (1.000000) | 3.07e-04 (1.000000) |

**Table S10.** Replication, the synthetic training set trajectory of each of the three independent executions of the identical optimization prompt, all ten iterations (seconds, 100 genotypes × 100 models). The prompt is the one used for the first optimization loop, so all three replicates optimized against the frozen synthetic training set. The three agents did not start from a common baseline. They made different decisions about how much of the starting script's genetics to repair before optimizing anything, so iteration 1 already differs 1.6-fold across replicates, and the replicate that folded its correctness rewrite into iteration 1 reports the smallest ratio at the end.

| **Iteration** | **Replicate A** | **Replicate B** | **Replicate C** |
|---|---:|---:|---:|
| v1 | 140.91 | 90.51 | 126.93 |
| v2 | 158.52 | 70.13 | 80.48 |
| v3 | 125.83 | 63.93 | 45.95 |
| v4 | 119.62 | 61.42 | 34.57 |
| v5 | 49.80 | 44.13 | 18.09 |
| v6 | 47.09 | 44.75 | 14.73 |
| v7 | 41.13 | 43.76 | 13.72 |
| v8 | 23.09 | 45.26 | 10.80 |
| v9 | 18.58 | 27.81 | 8.73 |
| v10 | 13.54 | 21.80 | 7.39 |

**Table S11.** Replication, every measured run on 1000G validation data. The table covers three replicate scorers plus the validated 1000G-trained scorer as an external reference, five shapes, three independent data draws per shape, and two executions per draw at the four small shapes and one at 100 × 100. That is 108 runs, of which 96 completed, 12 were killed out of memory and none timed out. Draws are identical across scorers, and replicate order within each shape and draw was rotated so that no scorer sat in the cold-page-cache slot more often than the others. Values are seconds, and where a cell holds two entries they are the two executions of the same input, in execution order.

| Shape | Draw | A | B | C | Reference |
|---|---|---|---|---|---|
| 1 × 1 | 1 | 38.9 / 39.0 | 13.5 / 12.0 | 28.2 / 28.0 | 4.1 / 4.0 |
| 1 × 1 | 2 | 39.0 / 39.0 | 15.0 / 15.0 | 40.0 / 40.0 | 5.0 / 5.0 |
| 1 × 1 | 3 | 39.0 / 40.0 | 16.0 / 16.0 | 39.0 / 40.0 | 5.0 / 5.0 |
| 1 × 10 | 1 | 41.0 / 43.0 | 16.0 / 16.0 | 28.0 / 28.0 | 5.0 / 6.0 |
| 1 × 10 | 2 | 43.0 / 40.0 | 16.0 / 16.0 | 40.0 / 41.0 | 6.0 / 6.0 |
| 1 × 10 | 3 | 42.0 / 42.0 | 17.0 / 17.0 | 40.0 / 39.0 | 5.0 / 5.0 |
| 10 × 1 | 1 | 72.0 / 77.1 | 26.0 / 32.0 | 53.0 / 53.0 | 11.0 / 11.0 |
| 10 × 1 | 2 | 60.0 / 57.0 | 24.0 / 30.0 | OOM / OOM | 12.0 / 11.0 |
| 10 × 1 | 3 | 69.0 / 63.0 | 25.0 / 31.0 | 51.0 / 51.0 | 11.0 / 11.0 |
| 10 × 10 | 1 | OOM / 78.3 | 30.0 / 31.0 | 50.9 / 54.0 | 12.4 / 13.0 |
| 10 × 10 | 2 | 69.0 / 67.0 | 46.0 / 37.0 | OOM / OOM | 18.0 / 17.0 |
| 10 × 10 | 3 | 72.0 / 84.0 | 29.0 / 37.0 | 53.0 / 50.0 | 13.0 / 11.0 |
| 100 × 100 | 1 | OOM | 154.1 | OOM | 151.1 |
| 100 × 100 | 2 | OOM | 149.1 | OOM | 123.1 |
| 100 × 100 | 3 | OOM | 178.1 | OOM | OOM |

**Table S12.** Replication, all twelve out-of-memory events, with the time survived and the origin of the kill. Out-of-memory results were recorded rather than retried. There are three regimes here. The first is deterministic (A and C at 100 × 100, every repeat), the second is data-driven (C at 10 × 10 and 10 × 1, only on draw 2, in both executions) and the third is execution-driven (A at 10 × 10 draw 1, which died on one execution and completed the identical input on the next).

| Shape | Scorer | Draw | Execution | Died after | Kill origin |
|---|---|---|---|---|---|
| 10 × 1 | C | 2 | 1 | 53.1s | host cgroup (6 processes killed) |
| 10 × 1 | C | 2 | 2 | 52.1s | host cgroup (6 processes killed) |
| 10 × 10 | C | 2 | 1 | 53.2s | host cgroup (6 processes killed) |
| 10 × 10 | C | 2 | 2 | 51.2s | host cgroup (6 processes killed) |
| 10 × 10 | A | 1 | 1 | 98.5s | host cgroup (2 processes killed) |
| 100 × 100 | C | 1 | 1 | 62.2s | host cgroup (24 processes killed) |
| 100 × 100 | C | 2 | 1 | 68.0s | host cgroup (22 processes killed) |
| 100 × 100 | C | 3 | 1 | 65.5s | host cgroup (14 processes killed) |
| 100 × 100 | A | 1 | 1 | 76.4s | host cgroup (28 processes killed) |
| 100 × 100 | A | 2 | 1 | 79.0s | host cgroup (24 processes killed) |
| 100 × 100 | A | 3 | 1 | 81.7s | host cgroup (30 processes killed) |
| 100 × 100 | Reference | 3 | 1 | 119.1s | GPU allocator (CuPy OutOfMemoryError) |

**Table S13.** Replication, variance components of $\log_{10}$ wall clock, as absolute variances and as percentages of the total, from a two-way random-effects decomposition over scorer $\times$ data draw with two executions per cell. Fold-spread is 10 raised to twice the corresponding standard deviation, so it reads directly as a multiplicative range. The draw term is a main effect, so it holds only the part of a draw's influence that is shared by every scorer, and a draw effect specific to one program appears in the scorer $\times$ draw term instead. The 100 $\times$ 100 shape has no estimable design, because only one of the three scorers completes it.

| Shape | Var(agent run) | Var(data draw) | Var(interaction) | Var(re-execution) | % agent | % draw | % interaction | % re-exec | Fold-spread agent/draw/re-exec |
|---|---|---|---|---|---|---|---|---|---|
| 10 $\times$ 10 | 0.0540 | 0.0000 | 0.0053 | n/a | 91.1 | 0.0 | 8.9 | n/a | 2.92 / 1.00 / n/a |
| 10 $\times$ 1 | 0.0701 | 0.0008 | 0.0000 | 0.0024 | 95.6 | 1.1 | 0.0 | 3.3 | 3.38 / 1.14 / 1.25 |
| 1 $\times$ 10 | 0.0468 | 0.0002 | 0.0025 | 0.0001 | 94.4 | 0.3 | 5.1 | 0.2 | 2.71 / 1.06 / 1.05 |
| 1 $\times$ 1 | 0.0556 | 0.0015 | 0.0018 | 0.0002 | 94.1 | 2.6 | 3.0 | 0.3 | 2.96 / 1.20 / 1.06 |
| 100 $\times$ 100 | not estimable | n/a | n/a | n/a | n/a | n/a | n/a | n/a | n/a |

**Table S14.** Replication, per-model concordance of each replicate against the validated 1000G-trained scorer at $10 \times 10$, computed per model column across samples. Models differ in scale by orders of magnitude, so a correlation pooled over the whole matrix would mostly measure scale agreement. Medians are over the data draws and executions in which both scorers completed. NRMSD is the root-mean-square deviation in units of that model's own standard deviation across the panel.

| Replicate | Median per-model r | Worst model r | Models with r > 0.999 | Median NRMSD | Worst NRMSD |
|---|---|---|---|---|---|
| A | 0.834 | 0.251 | 0% | 1.046 SD | 28.88 SD |
| B | 0.999 | 0.990 | 50% | 0.080 SD | 0.54 SD |
| C | 0.996 | 0.989 | 30% | 0.151 SD | 1.00 SD |

**Table S15.** Memory-bounded shape ladder on 1000G data. Every run went through a wrapper placing the job in a systemd scope with a 72 GB host memory cap and swap disabled, a 48 GB CuPy device limit, and a machine-wide free-memory watchdog. Peak memory is total memory minus the minimum machine-wide available memory, sampled twice a second across the whole process tree.

| Rung | Repeats | Wall clock | Peak machine memory | Model loci | Targets | Model rows | Model blocks |
|---|---|---|---|---|---|---|---|
| 100 × 100 | 3 | 171.5 s / 197.1 s / 192.8 s | 35.3 / 41.0 / 36.8 GB | 9.9 to 23.0 M | 10.1 to 18.7 M | 58.6 to 80.4 M | 1 |
| 500 × 500 | 1 | 1,461.4 s | 60.1 GB | 22.6 M | 19.2 M | 348.8 M | 2 |
| 1000 × 1000 | 1 | 3,471.1 s | 63.6 GB | 27.3 M | 21.7 M | 629.4 M | 4 |
| 3202 × 5385 (all) | 1 | 23,526.6 s (6.5 h) | 70.1 GB | 34.9 M | 27.7 M | 3,452.8 M | 35 |

**Table S16.** Per-stage timings at the two largest shapes for memory-bounded scorer.

| Stage | 1000 × 1000 | 3202 × 5385 (all) |
|---|---:|---:|
| Model locus pass (all scoring files) | 70.8 s | 397.0 s |
| VCF pass A, target union and ordinals | 1,580.3 s | 8,014.0 s |
| VCF pass B, dosage matrix | 1,519.7 s | 7,629.2 s |
| Model re-read (full decode, in blocks) | 74.9 s | 1,002.3 s |
| Variant matching | 24.4 s | 195.2 s |
| Weight matrix build | 15.7 s | 47.0 s |
| Sparse product (cuSPARSE) | 176.5 s | 6,216.0 s |
| Write output | 0.3 s | 4.7 s |
| Total | 3,471.1 s | 23,523.4 s |
| VCF ingestion share | 89.3% | 66.5% |
| Sparse product share | 5.1% | 26.4% |

**Table S17.** The non-additive defect, quantified over the full 3,202-sample panel. Exactly four of the 5,385 catalog models contain a dominant or recessive row, 21 rows in total, of which 16 matched a target in the full run, 16 out of 3.17 billion matched entries. The columns compare the repaired scorer against the inherited one, in which cuSPARSE's F-contiguous output made the non-additive kernel write into a discarded copy.

| **Model** | **Non-additive rows** | **Correct panel mean ± sd** | **Broken panel mean ± sd** | **Median retained** | **Median $\|\Delta\|$ (SD units)** | **Max $\|\Delta\|$ (SD units)** | **r (broken vs correct)** |
|---|---|---|---|---|---|---|---|
| PGS000318 | 2 of 2 rows recessive | -0.1335 ± 0.1079 | -0.1337 ± 0.1080 | 100% | 0.0000 (0.00) | 0.0021 (0.02) | 1.0000 |
| PGS000319 | 2 of 2 rows recessive | 0.2896 ± 0.1222 | 0.2882 ± 0.1223 | 100% | 0.0000 (0.00) | 0.0143 (0.12) | 0.9994 |
| PGS000802 | 7 dominant + 7 recessive of 19 rows | 2.1014 ± 0.4032 | 0.8524 ± 0.2675 | 41% | 1.2230 (3.03) | 2.5070 (6.22) | 0.6858 |
| PGS004700 | 3 of 12 rows recessive (none matched this panel) | 0.1499 ± 0.1920 | 0.1499 ± 0.1920 | 100% | 0.0000 (0.00) | 0.0000 (0.00) | 1.0000 |

**Table S18.** Full-catalog reference build and parameters. The build extends the scoring stage of the memory-bounded scorer and adds an output stage. Nothing upstream of scoring changed, and the raw scores are bit-identical to the inherited scorer's wherever a model is purely additive.

| **Parameter** | **Value** |
|---|---|
| Samples scored | 3,202 |
| Models scored | 5,385 |
| Target loci | 27,683,082 |
| of which SNP | 25,460,374 |
| Model loci read | 34,921,784 |
| Minimum locus call rate for S | 0.95 |
| Deployability threshold on $|S|/N$ | 0.75 |
| Minimum per-sample coverage of $|S|$ | 0.99 |
| Reference sample set | unrelated |
| Reference samples | 2,590 |
| Genotype PCs | 10 |
| PCA variants | 72,437 |
| PCA minor allele frequency floor | 0.05 |
| PCA variant spacing (bp) | 20,000 |
| Planning budget (GB) | 36.0 |
| Model blocks | 26 |
| Score tile (samples) | 15 |
| Models above the coverage threshold | 5,139 |

**Table S19.** Reference release, coverage, deployability and ancestry stratification across all 5,385 PGS Catalog models on the 1000 Genomes panel. Coverage is $|S|/N$, the fraction of a model's rows that both matched a target and were called in at least 95% of the reference samples. Super-population spread is the separation between the most and the least extreme super-population means, in units of the model's single pooled standard deviation. That is the number which says whether a pooled or an ancestry-adjusted percentile is the honest one for that model.

| **Quantity** | **Value** |
|---|---|
| Models in the release | 5,385 |
| Models with a defined reference distribution | 5,364 |
| Models matching nothing on this panel | 21 |
| Median model coverage $|S|/N$ | 0.943 |
| Models at coverage $\geq 0.75$ (flagged deployable) | 5,139 |
| Models below 0.75 (flagged, not dropped) | 246 |
| Median super-population mean spread | 1.11 pooled SD |
| Models with spread > 1 pooled SD | 58.5% |
| Models with spread > 2 pooled SD | 12.3% |
| Median score variance explained by 10 genotype PCs | 19.1% |
| Models with > 50% of variance explained by 10 PCs | 15.7% |
| Reference samples (unrelated) | 2,590 |
| Trio children scored but held out of the reference set | 612 |

[4] Emre Green, Xinmeng Liao, and Adil Mardinoglu. G-PGS: GPU-enabled whole-catalog PGS outperforms disease-specific PGS in complex disease profiling. Research Square, 2026.
